# mnDINO: Accurate and robust segmentation of micronuclei with vision transformer networks

**DOI:** 10.64898/2026.03.09.710648

**Authors:** Yifan Ren, Louise Morlot, J. Owen Andrews, Emil Peter Thrane Hertz, Niels Mailand, Juan C. Caicedo

## Abstract

Recent advances in cell segmentation successfully produce models that generalize across various cell-lines and imaging types. However, these methods still fail to recognize subcellular structures such as micronuclei (MN), which are rare and tiny DNA-containing structures found outside of the main nucleus and observable under the microscope. While they can be hard to recognize in images, studying MN formation is of great interest because of their relationship to chromosome instability, genotoxicity, and cancer progression. Here we present a segmentation model, mnDINO, to segment micronuclei in DNA stained images under diverse experimental conditions with very high efficiency and accuracy. To train this model, we collected a heterogeneous set of images with more than five thousand annotated micronuclei. Trained with this diverse resource, the mnDINO model improves the accuracy of MN segmentation, and exhibits strong generalization across microscopes and cell lines. The dataset, code, and pre-trained model are made publicly available to facilitate future research in MN biology.

## Introduction

Micronuclei (MN) —formed from lagging and acentric chromosome fragments that fail to become part of the daughter nucleus— are frequently found in tumorigenesis, and genotoxic stress conditions. The driving factors of MN formation are an active field of research, and include chromosome segregation errors caused by mitotic defects and DNA damage ^1–3^. The detection and quantification of MN serves as an indicator of genomic instability, because MN presence signals defects in mitosis, DNA replication or DNA repair. Moreover, MN have been linked to catastrophic events for the cell, probably via chromosomal rearrangements and chromothripsis ^4^. The study of MN biology is often conducted with imaging ^5^, which is the only tool that enables the observation of their emergence and phenotypic characteristics.

MN are typically quantified through observation of cells under a microscope, and can involve either manual scoring ^6,7^ or automated detection ^8^. Manual scoring is extremely time consuming and prone to fatigue and inter-scorer variability ^9^. Automated MN detection can be computationally challenging due to the small nature of the target objects, typically ranging from 1/16 to 1/3 of the mean diameter of nuclei ^10,11^. In addition, automated MN detection in images can be sensitive to noise, which results in artifacts —such as debris and background staining— being misclassified as true MN. The morphological features of MN (e.g., small size, varying intensity, proximity to other objects) are very unique and challenging for classical image segmentation methods.

Deep learning for cell segmentation has made great progress with models that identify cells in a wide range of microscopy images. Specifically, nucleus segmentation was first attempted across imaging experiments in the 2018 Data Science Bowl ^12^, showing potential in generalizing to any microscopy image. Whole-cell segmentation was approached next, and the Cellpose models ^13–15^ demonstrated high accuracy and strong generalization. Other methods such as StarDist ^16^, microSAM ^17^, and CellSAM ^18^, among others ^19^, have introduced additional innovations to make cell segmentation accurate and reliable. Importantly, these models have been trained with a large and diverse dataset of manually annotated images, primarily targeting cells and nuclei. Unfortunately, these models cannot identify other subcellular structures —such as MN— because of their assumptions about cell shapes and sizes, reflected in the annotations of datasets created for that purpose, and in the model design.

The successful approach used to segment cells in arbitrary microscopy images can in principle be extended to other cellular structures, including MN. However, there are two main problems with implementing models for MN segmentation. First, the lack of datasets with sufficiently diverse example images and high-quality manual annotations, which are challenging to obtain due to the rarity of spontaneous micronucleated cells and to the tiny size of MN. Second, the adaptation of models to accurately detect these objects, which are significantly smaller and more difficult to find than cells. Recently, these challenges were investigated with the MNFinder model ^8^, an ensemble of neural networks specialized in MN detection, achieving remarkable accuracy. Inspired by their work and the wider progress in cell segmentation, we aim to create a generalizable model to detect MN in DNA stained micrographs to investigate its mechanisms, as well as its effects in diverse biological problems.

## Results

### A heterogeneous dataset of annotated micronuclei

We curated a dataset of 232 DNA stained images of nuclei with 5,685 manually annotated micronuclei (MN) (Fig. 2). These images come from four experiments that involve different microscopes and varied experimental settings (Fig. 2b,c). MN are rare events under normal conditions, thus to increase their frequency in the collected images, all experiments used perturbations to induce MN formation. Two of the four experiments come from publicly available datasets that have been previously used to study nucleus and MN segmentation. These datasets are BBBC039 ^20,21^ and MNFinder_data ^8^, and involve U2OS and RPE1 cells treated with chemical perturbations. The other two experiments were conducted for this study, and we call them *mnDINO_data01* and *mnDINO_data02*. For these experiments, we used HeLa (mnDINO_data01 and mnDINO_data02) or RPE1 p53 -/- cells (mnDINO_data02) infected with a pooled library of distinct CRISPR-Cas9 guide RNAs targeting a diverse set of genes and genomic locations for CRISPR knockout (mnDINO_data01) or CRISPR interference (mnDINO_data02).

The four subsets were acquired with various microscopes using different objectives and magnifications (20X for BBBC039, MNFinder and mnDINO; 40X for MNFinder) and at different resolutions, resulting in varied numbers of cells per image and different object sizes. MN were manually annotated while nuclei were detected using Cellpose3 ^15^. We collected the size of nuclei and MN in all images and calculated their area in µm^2^ according to the camera parameters; their distributions show a large difference between nuclei and MN (Fig. 2d). The proportion of micronucleated cells is also different from experiment to experiment (Fig. 2e), with some experiments having more abundance of MN while others exhibit low occurrence. In addition, the distribution of MN sizes across experiments is also variable (Fig. 2f), which may be explained by differences in cell lines and perturbations. All these factors illustrate the challenges of MN detection, which is difficult because of the size of objects, their rare occurrence, and their morphological variations due to experimental conditions.

Our curated dataset brings together images with diverse cell lines, microscope settings, and perturbation types, resulting in a heterogeneous collection of annotated data to train models for MN segmentation. Following best practices in machine learning, we split the dataset in three subsets for training, validation, and test (Suppl. Tab. 1). The resulting heterogeneous dataset is a valuable asset to investigate MN segmentation models that generalize to different conditions.

### mnDINO accurately finds micronuclei in diverse images

Our model, mnDINO, is a ViT transformer backbone pre-trained on natural images with the DINOv2 algorithm. A segmentation head is attached to the backbone to transform local patch features into segmentation masks ^22,23^. The segmentation head is trained using supervised learning and the backbone is simultaneously fine-tuned to support the segmentation task. For model training, we use the 121 training images in our dataset containing 3,407 example MN objects. During training, patches of 256×256 pixels are randomly generated around regions where MN are visible, and we discard patches without MN. To make predictions, we run our model using a sliding window strategy to cover all areas of a high resolution DNA image. That is, patches of 256×256 pixels are evaluated by moving the window 32 pixels in the horizontal and vertical directions.

The results show that mnDINO has strong performance in all the test subsets of the heterogeneous dataset (Fig. 3a). Performance is measured using object-centric precision and recall scores, assuming an object is correctly detected if the prediction mask has an intersection-over-union overlap of at least 0.1, following previously proposed evaluation protocols for MN ^8^. mnDINO achieves 75% precision on average across all subsets indicating a general ability to identify real objects in diverse imaging conditions. Our model also achieves 82% recall on average, which indicates that most of the real MN objects are detected. These results are in contrast to MNFinder, which is the best baseline approach specialized in MN segmentation, and obtains 65% precision and 77% recall, on average. That is, our model can improve performance by 15% in precision and 6% in recall.

We also compare the performance of mnDINO against other methods that have been designed for general purpose cell segmentation. The pre-trained microSAM ^17^ model achieved 22% precision and 3% recall on average without fine-tuning, which illustrates how difficult it is to segment MN in DNA stained micrographs. Fine-tuning microSAM to segment MN did not result in better performance. The Cellpose3 nuclei model was more responsive to fine-tuning and achieved 50% precision and 18% recall on average, which is still far from satisfactory. Both microSAM and Cellpose were designed and pre-trained to segment full cells, which are at least one order of magnitude bigger than MN. Their low performance may be explained by their strengths at segmenting objects at the cellular scale rather than at subcellular resolution.

### Predicted objects closely match original objects

Qualitative inspection of segmented MN shows that mnDINO correctly finds their location and covers most of their spatial extent (Fig. 4). MN sizes vary greatly depending on a combination of experimental factors, including cell line, perturbation type, and imaging parameters; MN detection depends on the correct interpretation of the image context. The mnDINO model –trained on a heterogeneous dataset that covers diverse conditions– is able to successfully segment MN in held-out test images (Fig. 4, third column). We observe that the pre-trained model MNFinder does a good job at identifying MN in most cases, but can introduce artifacts and false positives when nucleus morphology is complex (Fig. 4, fourth column). Cellpose and microSAM miss the majority of the MN objects, especially when they are very small (Fig. 4 columns 5 and 6). These qualitative results illustrate how difficult MN segmentation can be.

The extremely small size of MN is what makes their segmentation challenging. We analyze the size of predicted masks produced by mnDINO to determine any deviations from the real size of manually annotated MN (Fig. 3b). The results show that our model is capable of producing masks that have similar size to the original objects in test images. For most predictions, mnDINO underestimates the size of MN and produces masks that have less pixels than the real objects (points below the gray line in Fig. 3b). However, this offset is not too large, the correlation is very high (0.8733 Pearson correlation and 0.7627 R^2 coefficient), and for practical applications the actual detection may be more important than the exact number of identified pixels.

mnDINO obtains accurate segmentation results without the need to adjust input image sizes. Adjusting the size of the images to a common scale did not work better and sometimes performed worse (data not shown). Our model can correctly identify MN under drastic magnification changes (e.g., 20X vs 40X) because of two aspects of our training strategy: first, a heterogeneous dataset which offers learning potential from large biological and technical variation. (2) Data augmentations that randomly simulate smaller or bigger objects. Given that the model is trained to segment both the nucleus and MN, it learns to segment objects that are small relative to the nucleus size. Therefore, mnDINO learned to pay attention to the context to correctly segment MN.

### mnDINO generalizes to microscopes and cell lines

The mnDINO model was trained with a heterogeneous dataset that includes images from diverse experimental conditions. We sought to determine if the model is capable of generalizing to out-of-distribution images by conducting two evaluations. The first evaluation aims to reveal how sensitive the model is to different microscopes and camera specifications, while the second evaluation aims to measure performance changes when the cell line is different from those used in the training set. We systematically conducted cross-validation evaluations by leaving out training images according to the variable of interest. The results show that the performance of mnDINO drops slightly because of variations in the validation images not observed during training (Fig. 5). Nevertheless, overall performance remains high, which indicates that the majority of micronuclei are still correctly detected.

Generalization to new microscopes was simulated by leaving out images from the four subsets in a cross validation experiment. Each subset has a different combination of microscope equipment and digital camera, resulting in optical variations, and different pixel sizes and image resolutions. We observe that the hold-out models maintain similar performance to the model trained with the full dataset, dropping only by 2.6% on average. Note that mnDINO automatically adapts to differences in image sizes or pixel resolution, and does not require any manual adjustments. A similar cross-validation evaluation was conducted by isolating cell lines in the training set, and the observed drop in performance is slightly higher at 8.2% on average. The gap was larger for U2OS (∼12%) and HeLa (∼8%) cells, suggesting that MN detection is sensitive to morphological differences across cell lines, such as nucleus phenotypes or MN with varying shapes / sizes. For reference, the performance change when generalizing to RPE1 cells was only 2.4% in F1-score.

The cross validation experiments reveal how sensitive the model is to out of distribution data, which is an estimation of model performance when used without re-training or fine-tuning. Note that even when trained without data specific to microscope or cell line, the model still performs better than Cellpose or microSAM on the MN segmentation problem (Figs. 3 and 5). Performance is also better than MNFinder in these generalization tasks, especially for the most challenging cell-line generalization problem. Overall, the results show that mnDINO maintains high performance when tested on out-of-distribution images, exhibits robustness to microscope variations, and can be used across cell lines with minor drops in performance.

### mnDINO is computationally efficient

Our model predicts segmentation masks in overlapping regions of 256×256 pixels, following a sliding window approach with variable step size (Fig. 1). The smaller the step size, the more redundant predictions are, resulting in potentially better performance and higher computational cost. We evaluated the impact of step size on model performance (Fig. 6a), and found that the F1 score can improve between 5% and 17% when the step size changes from no overlap (256 pixels) to high overlap (16 pixels). While the gains can be significant, the cost of processing an image of 1024×1024 pixels grows exponentially from less than a second with no overlap to more than a minute with high overlap (Fig. 6b). Compute time was measured on an NVidia A100 GPU. Given the tradeoff between performance and compute time, the step size parameter needs to be adjusted depending on the compute budget and required accuracy. In all our experiments, we used a step size of 32 pixels, which maintains high performance and results in an approximate segmentation time of 25 seconds for images of 1024×1024.

**Figure 1.**
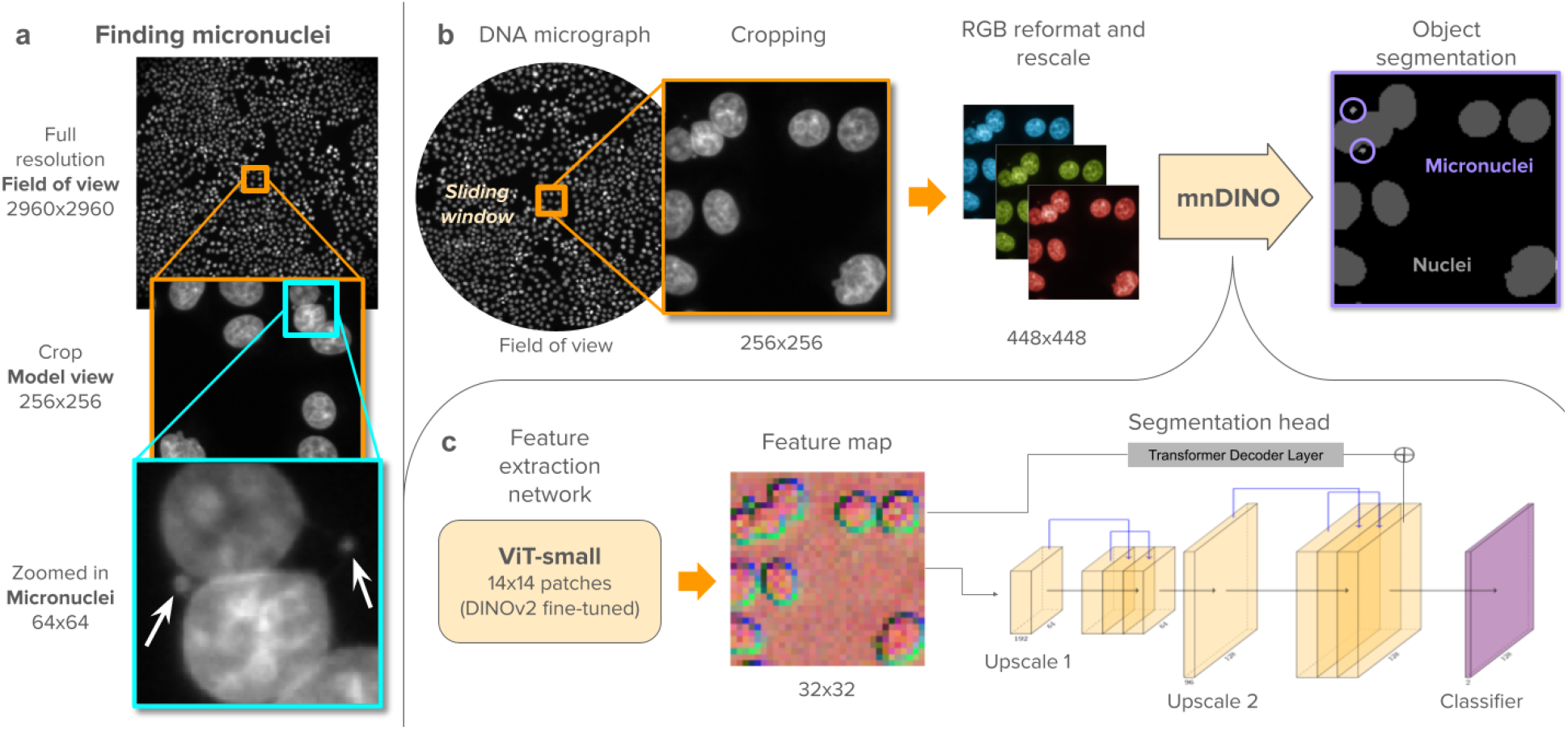
Finding micronuclei in DNA-stained micrographs. a) Micronuclei are rare and very tiny, requiring extreme zoom in to be seen by the human eye. Top: full resolution field-of-view at 20X magnification with ∼1,000 visible nuclei. Middle: an image crop of the size seen by the model with ∼10 nuclei. Bottom: an extreme zoom-in region showing 2 micronuclei, which can occupy less than 20 pixels of area. b) Proposed approach: images are cropped into 256×256 regions using a sliding window. Crops are transformed from grayscale to RGB and interpolated into 448×448 pixels. The interpolated crops are processed by the mnDINO model, which produces masks for both nuclei and micronuclei. c) Architecture of mnDINO: a fine-tuned ViT backbone encoder generates local patch features, which are processed by a segmentation head composed of pixel and transformer decoders.

**Figure 2.**
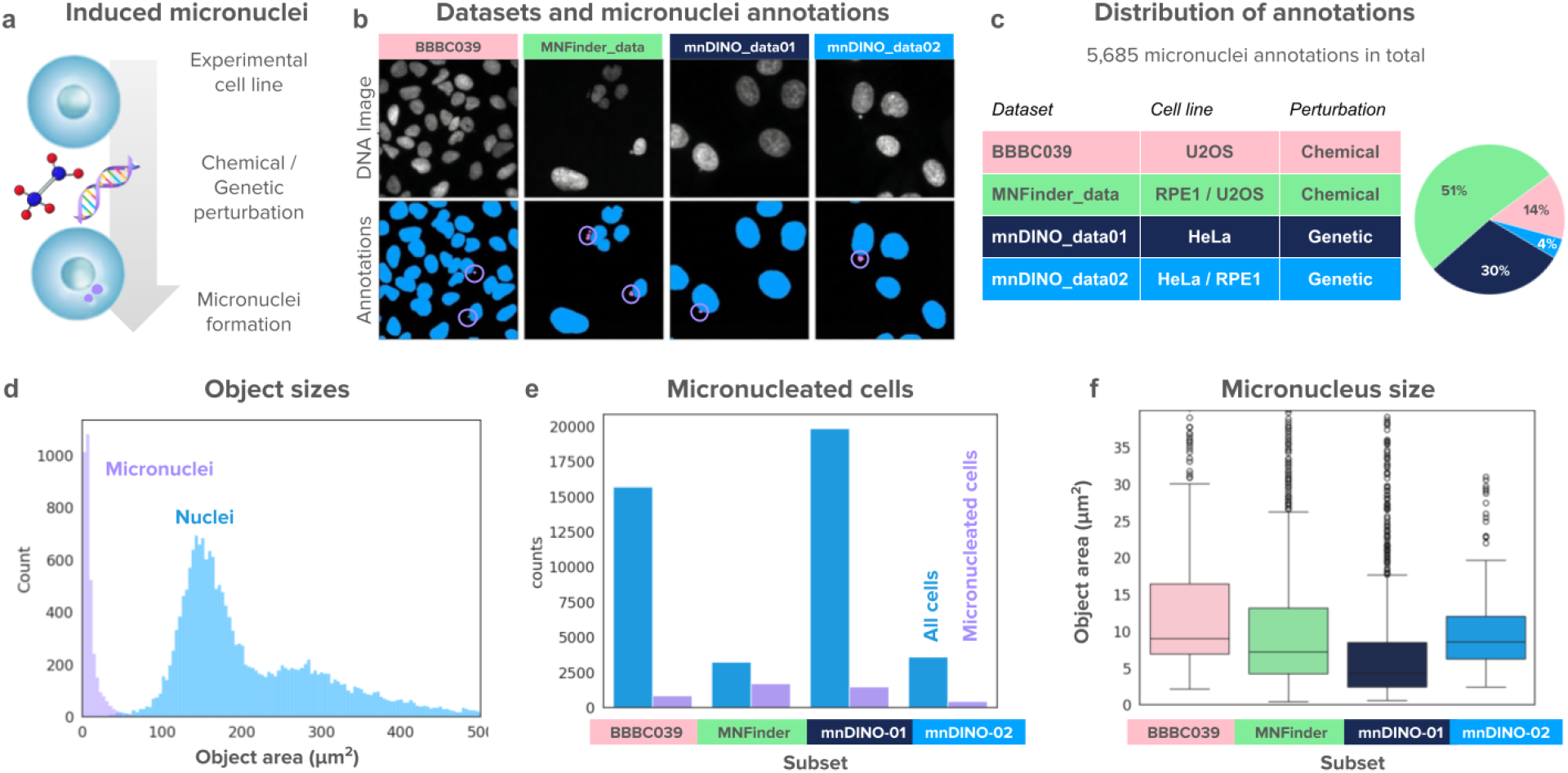
Annotated MN image dataset. a) Chemical or genetic perturbations are used to induce micronuclei formation on experimental cell lines. b) Example images from each of the four data sources. Top row: DNA stained images, bottom row: segmentation masks (nuclei in blue and MN in purple). c) Experimental properties of the four datasets and pie chart of the distribution of MN annotations across the datasets. d) Distribution of object sizes in all images of the dataset: horizontal axis is object area in µm^2^ and vertical axis is frequency. e) Cell counts per image subsets: blue bars are all cells, purple bars are micronucleated cells. f) Distribution of MN sizes in each subset: vertical axis in µm^2^.

**Figure 3.**
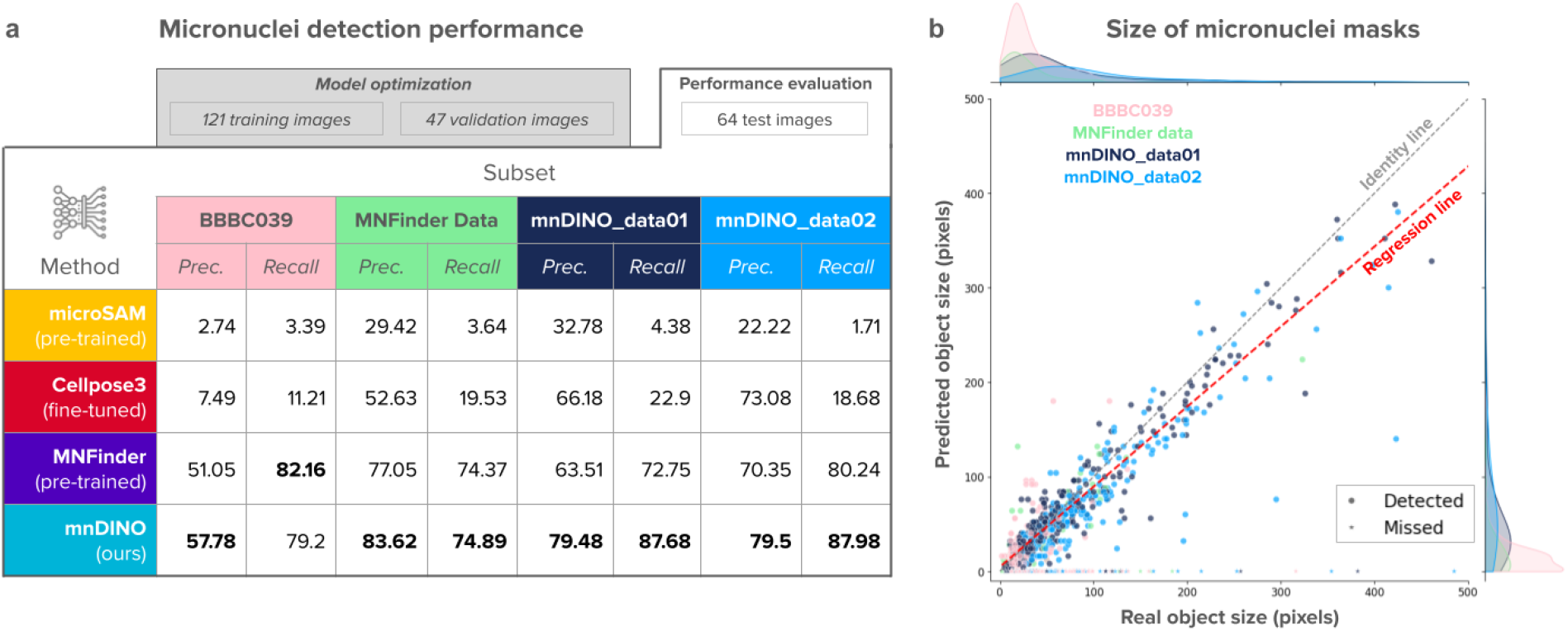
Quantitative evaluation of MN segmentation. a) Evaluation results comparing precision and recall performance of four models (rows) on four subsets of test data (columns). Higher scores indicate better performance. Bold numbers indicate the best performance across columns. b) Comparison between real object sizes (horizontal axis) and size of objects predicted by mnDINO (vertical axis). Each point is one of the 1,238 annotated MN in the test images across the four subsets (colors: pink: BBBC039, green: MNFinder_data, dark blue: mnDINO_data01, light blue: mnDINO_data02). Detected objects are circles, missed objects are small stars. Point colors indicate source subset. The gray line indicates the theoretical identity line where real and predicted sizes are the same. The red line indicates the observed regression line calculated across all validation data points (from all subsets), which has a Pearson correlation of 0.8733 and R^2 coefficient 0.7627.

**Figure 4.**
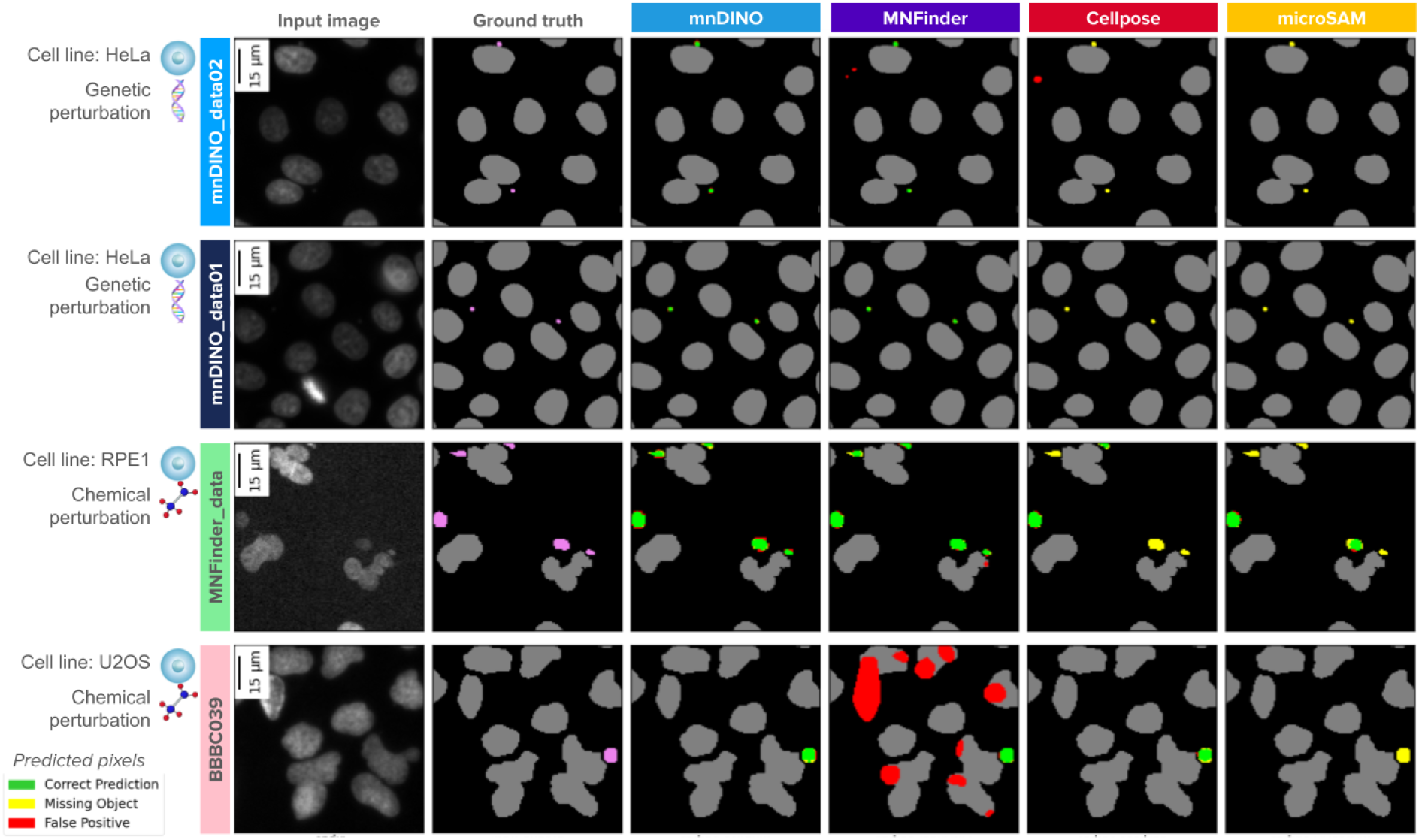
Visualization of micronuclei segmentations. Qualitative illustration of MN segmentations with one example image from each test subset (rows) and segmentation results by different methods (columns). The information at the left of the grid indicates the cell line and type of perturbation of the example image (first column). All example images are scaled to the same pixel size. The ground truth image (second column) shows nucleus masks in gray and MN masks in magenta. The segmentations produced by methods (third to sixth columns) follow the color convention for predicted pixels: green for correct, yellow for missing, and red for false positives. The gray nucleus masks from the ground truth are preserved in gray for reference.

**Figure 5.**
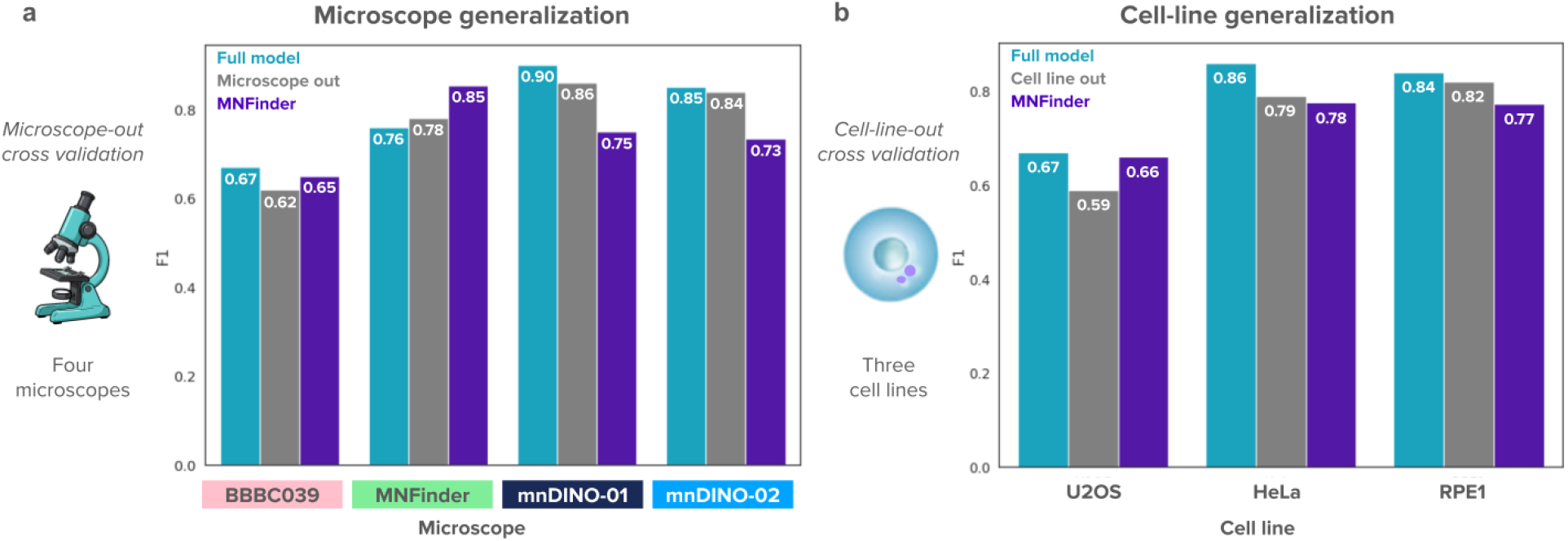
Model generalization to out-of-distribution conditions. a) Performance evaluation on the validation set when the training set does not contain any images from the indicated study. The horizontal axis lists the four validation subsets; each used a different microscope for image acquisition. The vertical axis reports performance on the segmentation task using the F1 score, which is the harmonic mean of precision and recall. The plot compares performance of the full model (blue bars, trained with all images) against models trained leaving one microscope out (gray bars, trained without images from the indicated study), and the pretrained MNFinder model (purple bars, not retrained on the held-out subsets). b) Performance evaluation of mnDINO when the training set does not contain any images from the indicated cell line. The horizontal axis lists the three cell lines and the vertical axis reports performance using the F1 score. The plot compares the full model (blue bars) against models trained leaving one cell line out (gray bars, trained without images from the indicated cell line), and the pretrained MNFinder model (purple bars, not retrained on the held-out subsets).

**Figure 6.**
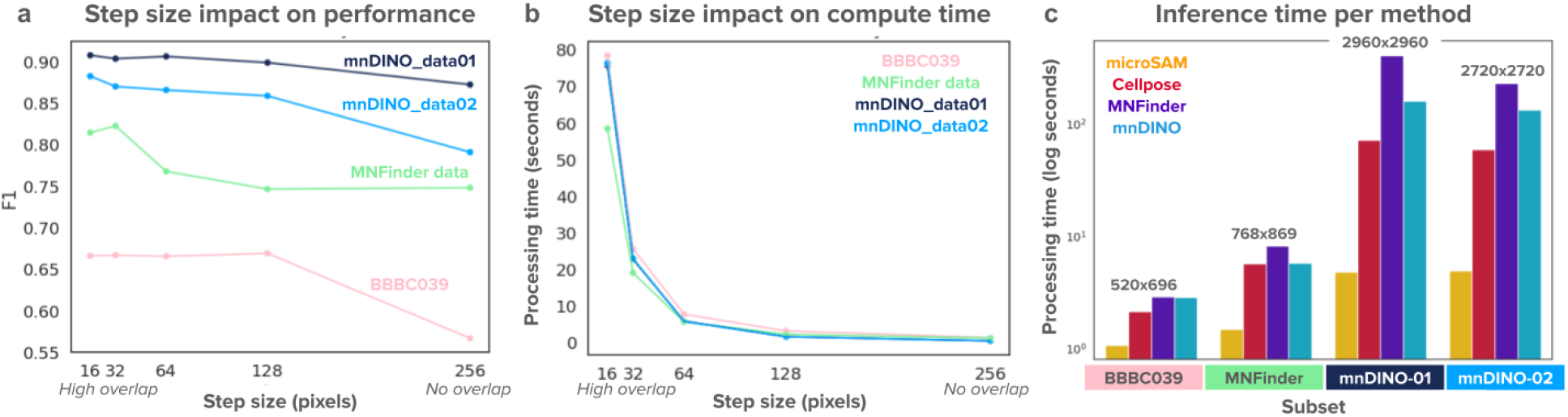
Computational cost of mnDINO. Step size is the parameter of mnDINO that controls the distance that the sliding window takes to make predictions in a high-resolution image (see Fig. 1). a) F1 score (vertical axis) as a function of step size (horizontal axis) evaluated in the validation sets of the four imaging subsets (colors). b) Inference time in seconds (vertical axis) as a function of step size (horizontal axis) for validation images in the four subsets (colors). Due to differences in image sizes across the subsets, the plot has been adjusted to report the time to process an image of constant area = 1024×1024 pixels. c) Comparison of inference time taken by different segmentation methods on the four subsets. The validation sets are in the horizontal axis, and the log-inference time (seconds) is reported in the vertical axis. The bars are colored according to the segmentation method. Reported time is not normalized, indicating that larger images take longer times. Time reported with mnDINO is based on step size = 32 pixels.

The images in the four subsets evaluated in our experiments have widely different sizes, from 520×696 in the BBBC039 subset to 2,960×2,960 in the mnDINO_data01 subset. When comparing the processing time of image segmentation in all subsets, we found that the cost of running mnDINO is higher than MicroSAM and Cellpose, and lower than MNFinder (Fig. 6c). All methods implement redundant computations such as sliding windows and, in the case of MNFinder, ensemble networks. The cost for all methods is comparable and grows with image size; the differences may only be important at large scale when processing thousands of images. However, their segmentation performance is very different and results in different accuracies (Fig. 3). Therefore, we conclude that mnDINO is as efficient or more efficient than other methods while providing the best segmentation quality.

## Discussion

In this work, we presented mnDINO, a ViT-based model for accurate MN segmentation. The size and morphology characteristics of MN make this problem challenging for traditional cell segmentation methods, and require specialized data to train effective models. An important contribution of this work was the curation of a heterogeneous dataset with over five thousand manually annotated MN masks for images acquired under four different settings and three different cell lines. The diversity of the training data is one of the strengths of mnDINO, which enables model generalization to different technical and biological conditions. The resulting pre-trained model is accurate and efficient, and can be reused for MN detection in DNA stained micrographs to study MN biology in large scale studies. Our model is especially good at generalising across acquisition modalities (microscope, objective, magnification and resolution), making it available to the community with little to no retraining.

We compared the proposed model against existing methods for cell and MN segmentation, and found that mnDINO significantly improves performance while maintaining comparable computational efficiency. Unlike the baseline models, we do not introduce a new neural network architecture or algorithm in this work. Instead, we leverage generic building blocks from computer vision, such as the pre-trained, transformer-based feature extractor DINOv2, and a lightweight state-of-the-art segmentation head trained with standard supervised loss functions. The only domain-specific adaptation introduced in our method is the resizing of images to double the size for feature extraction to magnify small objects. The simplicity of our approach, combined with heterogeneous training data, results in accurate and efficient MN segmentation.

Cell segmentation has improved with foundation models trained with heterogeneous and multimodal microscopy images. While these methods now fuel the analysis of microscopy images for biological research, there is still a long way to go when it comes to identifying and understanding subcellular structures. These are more complex to segment than cells because of their small, thin, rare, and delicate morphologies. Yet, as microscopists and biologists make progress with experimental techniques to observe cells at high resolution and high throughput, we will need the ability to accurately quantify such subcellular structures. MN are an example of a rare and very difficult to detect phenotype and we show that they can be accurately segmented with a combination of the right data and the right models. mnDINO can be readily applied to large scale investigations of MN formation, and our proposed dataset can be used to bring MN detection to models that recognize many more subcellular structures in the future.

## Acknowledgements

The authors thank Paul Blainey for his invaluable advice and support to this project. We also thank Vidit Agrawal for supporting early data analysis.

## Author contributions

Conceptualization and Design: Y.R., J.O.A., E.P.T.H., J.C.C. Data Collection & Experiments: Y.R., L.M., J.O.A., E.P.T.H. Data Curation & Annotation: Y.R., L.M., E.P.T.H. Computational Modeling & Software: Y.R., J.C.C. Experiments & Data Analysis: Y.R., J.C.C. Writing and Reviewing: Y.R., L.M., J.O.A., E.P.T.H., N.M, J.C.C.

## Methods

### Datasets

To train, validate and test mnDINO, we produced and curated 232 images coming from diverse cell lines and microscopes. BBBC039 subset is cultured on U2OS cells, imaged on a 20× ImageXpress Micro epifluorescent microscope ^24^. We filtered out the nuclei by removing objects that are larger than 100 pixels, and those that are in contact with the edges of the images. Blank images are discarded after pre-processing, resulting in 159 images with 1,013 annotated micronuclei. Image size in this subset is 520×696 pixels. MNFinder data was obtained from their public repository ^8^, and we follow their train-validation-test split for model training and evaluation. All images are 768×869 pixels in size and display RPE1, U2OS, HeLa, and HFF cells. MNFinder images are taken from both Leica DMi8 laser scanning confocal microscope and Leica DMi8 with Adaptive Focus widefield microscope at 20× and 40× resolutions. mnDINO data subset 01 is generated from a Nikon Ti2 Eclipse inverted epifluorescence microscope, equipped with a IRIS9 camera, a 20X objective (Plan Apochromat, NA 0.8) and a 0.7X demagnifying C-Mount adaptor using HeLa cells, resulting in 18 high-resolution images of 2960×2960 pixels. 1,688 ground truth objects were manually annotated. mnDINO data subset 02 comes from a Nikon Ti2 Eclipse epifluorescence microscope, equipped with a Kinetix camera and a 20× objective (Plan Apochromat, NA 0.8) on HeLa and RPE1 p53-/- cells, where 6 high-resolution images at 2720×2720 pixels were annotated with 582 objects.

### Cell Pool Generation for mnDINO subset data

Cell pools with libraries of CRISPR/Cas9 guide RNAs were produced following the protocol in ^25^. Briefly, oligo pools containing gRNAs sequences were ordered from Agilent, and cloned into the CROPseq-Guide-Puro vector ^26^. Guide RNAs were selected to target a variety of genes and chromosomal locations. Plasmid libraries were transfected into 293Ts to be packaged into lentiviral particles. Cell lines were transduced with the lentivirus at an MOI of 0.1, and selected for successful integration with Puromycin.

For mnDINO data subset 01, the guide RNA library was designed to target the coding sequence of target genes for efficient CRISPR/Cas9 knockout, and was transduced into a HeLa cell line expressing wild type Cas9. For mnDINO data subset 02, the guide RNA library was designed to target the transcription start site of target genes for efficient CRISPR interference, and was transduced into HeLa and RPE1 p53-/- cell lines expressing *ZIM3* KRAB-dCas9 ^27^.

### Manual annotations

MNFinder cells are labeled with DAPI and H2B-FP, BBBC039 cells are stained with Hoechst and both mnDINO datasets with DAPI. mnDINO data annotation was created by experts who manually labelled the objects of interest using image editing software. Specifically, observers were asked to follow the 2018 Data Science Bowl procedure for cell mask annotations ^12^, which uses GIMP software to delineate each micronucleus in the images. Nuclei masks are generated using Cellpose3 ^15^, and we post-process nuclei masks by removing the pixels that overlap with the micronuclei masks.

### Model Architecture

Recent advances in self supervision and vision transformers, such as DINO (DIstillation with NO labels), have shown superior performance when compared to supervised and weakly supervised learning methods ^28^. We use DINOv2 as our model backbone to extract meaningful features from the cellular images ^29^, which is built upon the vision transformer that patchifies the input images into small tokens of 14×14 pixels, with an additional class token appended to aggregate all local information ^30^. DINO leverages the knowledge distillation by training a student network g_s_ to match the output of a teacher network g_t_. During training, the student network receives both global and local views of the image, and the teacher network only receives global views. The teacher parameters are updated through the exponential moving average of the student parameters. We use the DINOv2 ViT small backbone, and built a lightweight version of the state-of-the-art segmentation head Mask2Former ^22^.

Our implementation takes an image of 256×256 pixels as input and rescales it to 448×448 pixels. From this, DINOv2 generates a feature map of 32×32 tokens each with 384 features. The segmentation head reduces the extracted features by half and expands the spatial resolution by double using convolutional layers as follows: the input feature map is transformed from 32×32×384 to 64×64×192 in the first upscale block, then processed at that resolution by 3 convolutional layers. Next, the feature map is then transformed again to 128×128×96 in the second upscale block, and processed by an additional group of 3 convolutional layers. An additional embedding from the transformer decoder layer of 64×64×192 is interpolated to 128×128×192 and linearly projected into 128×128×96. This embedding is added into the output of the last convolutional layer. Finally, the combined features are passed into a classification head that produces both nucleus and micronucleus predictions in a 128×128×2 probability map. This probability map is interpolated to 256×256×2 to produce the final output. This model is used in a sliding window manner with a configurable step size to cover all the area of the original image.

### Model Training

In each of the four datasets, we randomly split 70% into training images, and 15% for validation and test images respectively, resulting in 121 training images, 47 validation images and 64 test images (Suppl. Tab. 1). The gray-scale images are randomly cropped around micronuclei by sampling the height and width from a Gaussian distribution N(256, 20) and then resizing into 256×256 pixels. This results in a data augmentation function that simulates varying object sizes. Other standard data augmentations are applied, including random rotations, horizontal flips, brightness and contrast adjustments. Then the crops are replicated into RGB channels to be fed to the backbone. We observed that the random sampling of crop resolutions help the model to be distortion-agnostic, while other augmentations improve model robustness.

For the loss function, we follow the 20:1 ratio of focal loss to dice loss that is reported in the Segment Anything Model ^23^. A customized dice loss is applied by forcing 0.8 weight to the micronucleus class and 0.2 weight to the nucleus class. The model is trained on a single NVIDIA A100 GPU for roughly 1.8 hours, we use AdamW optimizer (*β*_1_=0.9, *β*_2_=0.999) with learning rate 0.00001 and weight decay 0.000001 for 20 iterations ^31^, a cosine scheduler (*T*_max_=20, *η*_min_=0.000001) is also applied ^32^.

During training and validation, we observed a performance trade off between precision and recall (Suppl. Fig. 1 and 2). That is, when a model improves precision, recall decreases, and the other way around. This trade-off is the result of many factors, especially the small size of the objects. When the model gets real MN more accurately (improved precision), it may miss borderline instances, and when it covers most target objects correctly (improved recall), it may introduce false detections. After conducting rigorous cross-validation, we selected a model that favors high precision while maintaining a satisfactory level of recall.

### Segmentation Algorithm

Images need to be normalized using min-max pixel normalization (instance normalization) prior to making predictions, due to mnDINO being trained on normalized micrographs. A sliding window of 256×256 scans vertically and horizontally by a fixed step size, for every 4 crops collected we put them in a single batch to generate predicted probabilities, and average them across each row. We report the best performance of mnDINO on the test set using 32 step size. To test the generalizability of mnDINO, specific subsets that come from the same cell-line or microscope are held out during the training stage in a cross validation scheme. Then we evaluate the model predictions on the validation set using step size 64.

### Evaluation of performance

To evaluate the correctness of predicted micronuclei, we use the standard metric intersection-over-union (IoU) to identify if there are any pixel-wise matches between ground truth *T* and the predicted objects *E*. IoU is calculated as follows:

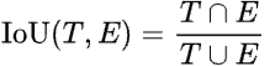

Given the IoU matrix among all predicted and ground truth objects, we use a 0.1 IoU threshold as a cutoff to select the matched objects ^8^. We then calculate the values of the confusion matrix: True Positives (TP), False Positives (FP), and False Negatives (FP), to calculate the standard performance metrics precision, recall and F1-score as follows:

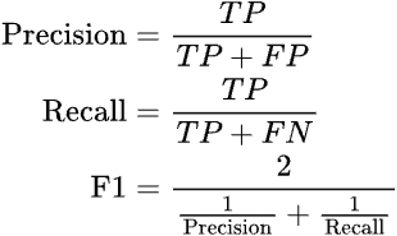

When evaluating Cellpose, we fine-tuned the model with all the available training data and adjusted learning rate, weight decay, epochs and minimum training masks, to improve accuracy. We made several attempts to fine-tune the microSAM model following available instructions and documentation; however, performance did not improve beyond what is achieved with the pre-trained model. We did not re-train or fine-tune the MNFinder model due to the two-stage model architecture (a detector and a segmentation model). In addition to being two-stage, it also has multiple model replicates to create an ensemble, which improves accuracy, but also increases computational complexity. Therefore, MNFinder was also evaluated with its publicly available pre-trained weights.

## Data availability

The annotated MN dataset, which contains the four pre-processed and integrated subsets, is publicly available on BioImage Archive ^33^: https://www.ebi.ac.uk/biostudies/bioimages/studies/S-BIAD2809. The BBBC039 dataset is available at https://bbbc.broadinstitute.org/BBBC039, and MNFinder data is available on Github: https://github.com/hatch-lab/mnfinder/tree/main/src/mnfinder. A metadata.csv file is also provided in annotated_mn_datasets that describes the magnification, pixel size, and cell line information.

## Code availability

The code needed to reproduce the experiments in this study is publicly available on Github: https://github.com/CaicedoLab/micronuclei-detection. The pre-trained mnDINO model is also publicly available on Hugging Face: https://huggingface.co/CaicedoLab/mnDINO.

