## Supplementary Materials for "mnDINO: Accurate and robust segmentation of micronuclei with vision transformer networks"

### 1. Dataset organization

This dataset was created by combining images from four sources, resulting in heterogeneous imaging devices and cell-lines. Here we present a quantitative description of the dataset, which is organized in four subsets, and was split in training, validation, and test partitions.

| Subset | Total | | Training | | Validation | | Test | |
|---|---|---|---|---|---|---|---|---|
| | Images | Objects | Images | Objects | Images | Objects | Images | Objects |
| BBBC029 | 159 | 1013 | 77 | 488 | 39 | 270 | 43 | 255 |
| MNFinder | 49 | 2402 | 30 | 1749 | 3 | 180 | 16 | 473 |
| mnDINO_01 | 18 | 1688 | 12 | 1021 | 3 | 345 | 3 | 322 |
| mnDINO_02 | 6 | 582 | 2 | 149 | 2 | 245 | 2 | 188 |
| **Total** | **232** | **5685** | **121** | **3407** | **47** | **1040** | **64** | **1238** |

**Supplementary Table 1**: Quantitative summary of the number of images and objects (micronucleus masks) in each dataset and train/validation/test subset. Rows are the four main subsets in the micronuclei dataset, and rows are organized by dataset split. The number of objects refers to the number of manually annotated micronucleus masks available.

### 2. Variance of Model Performance

We observed that retraining mnDINO results in some performance variation and a trade-off between precision and recall. To characterize this variation, we replicated the training process 10 times with the same best configuration (see Model Training in Methods) found during the hyperparameter tuning phase, and measured performance in the validation set. Supplementary Figure 1 reports the variation in precision and recall using box plots for each subset. The results indicate that performance can vary significantly from experiment to experiment, from very high to very low performance. Interestingly, when precision improves recall tends to deteriorate and the other way around, indicating a trade-off between both metrics. The trade-off between precision and recall is better observed in Supplementary Figure 2. The main reason for this trade-off is the small and challenging nature of micronucleus instances, which can be confused with noise, artifacts, and debris in the micrographs. From these replicate models, we chose a final model that has high precision (>80%) and comparable performance in recall (~80%), and use it for all the analysis in the main text.

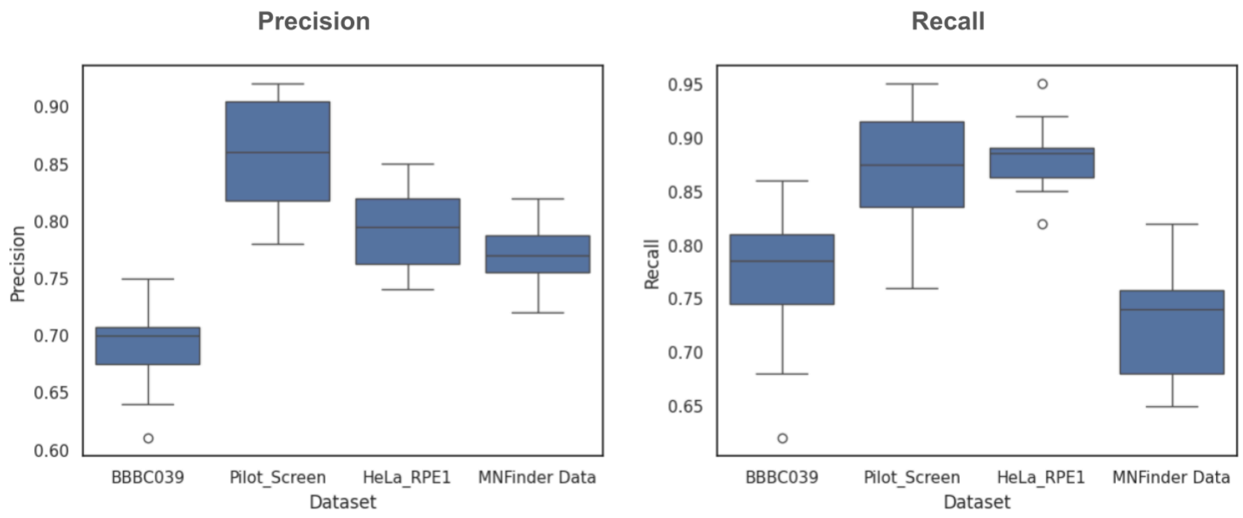


**Supplementary Figure 1**: Analysis of mnDINO model training variance. We repeat the training process 10 times to observe variations in precision and recall for each subset. The horizontal axis represents the validation data in the four subsets in our analysis. The vertical axis represents precision (left plot) or recall (right plot). In the box plots, the middle bar indicates the median, the box shows the first and third quartiles, and the dots represent outliers.

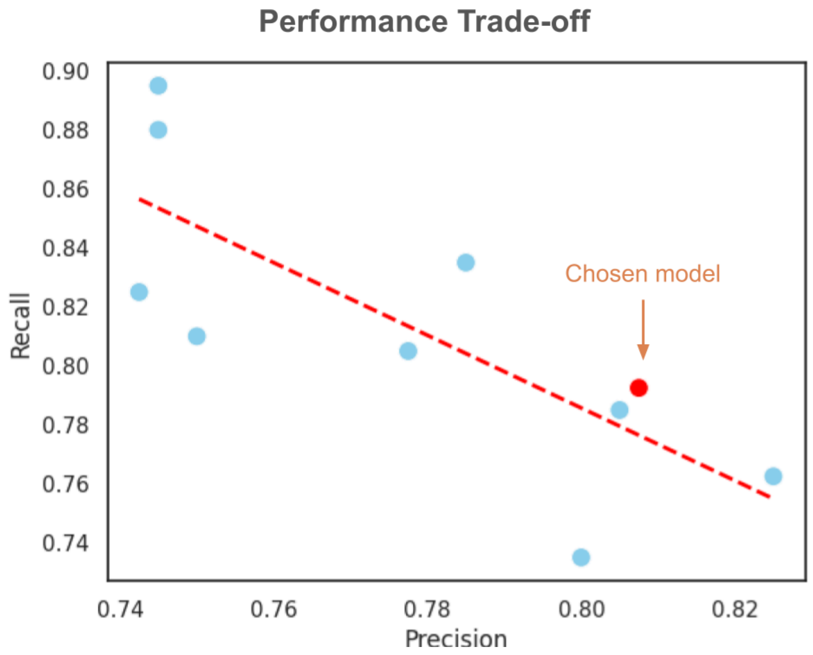


**Supplementary Figure 2**: Trade-off between precision and recall scores**.** The horizontal axis is precision and the vertical axis is recall. Each point represents one repetition of the training process with the same hyperparameters. The trend line has a negative Pearson correlation of -0.773. The chosen model is indicated in red.
